# Vector genetics, insecticide resistance and gene drives: an agent-based modeling approach to evaluate malaria transmission and elimination

**DOI:** 10.1101/2020.01.27.920421

**Authors:** Prashanth Selvaraj, Edward A. Wenger, Daniel Bridenbecker, Nikolai Windbichler, Jonathan R. Russell, Jaline Gerardin, Caitlin A. Bever, Milen Nikolov

## Abstract

Vector control has been a key component in the fight against malaria for decades, and chemical insecticides are critical to the success of vector control programs worldwide. However, increasing resistance to insecticides threatens to undermine these efforts. Understanding the evolution and propagation of resistance is thus imperative to mitigating loss of intervention effectiveness. Additionally, accelerated research and development of new tools that can be deployed alongside existing vector control strategies is key to eradicating malaria in the near future. Methods such as gene drives that aim to genetically modify large mosquito populations in the wild to either render them refractory to malaria or impair their reproduction may prove invaluable tools. Mathematical models of gene flow in populations, which is the transfer of genetic information from one population to another through migration, can offer invaluable insight into the behavior and potential impact of gene drives as well as the spread of insecticide resistance in the wild. Here, we present the first multi-locus, agent-based model of vector genetics that accounts for mutations and a many-to-many mapping cardinality of genotypes to phenotypes to investigate gene flow, and the propagation of gene drives in Anopheline populations. This model is embedded within a large scale individual-based model of malaria transmission representative of a high burden, high transmission setting characteristic of the Sahel. Results are presented for the selection of insecticide-resistant vectors and the spread of resistance through repeated deployment of insecticide treated nets (ITNs), in addition to scenarios where gene drives act in concert with existing vector control tools such as ITNs. The roles of seasonality, spatial distribution of vector habitat and feed sites, and existing vector control in propagating alleles that confer phenotypic traits via gene drives that result in reduced transmission are explored. The ability to model a spectrum of vector species with different genotypes and phenotypes in the context of malaria transmission allows us to test deployment strategies for existing interventions that reduce the deleterious effects of resistance and allows exploration of the impact of new tools being proposed or developed.

**Author summary:** Vector control interventions are essential to the success of global malaria control and elimination efforts but increasing insecticide resistance worldwide threatens to derail these efforts. Releasing genetically modified mosquitoes that use gene drives to pass on desired genes and their associated phenotypic traits to the entire population within a few generations has been proposed to address resistance and other issues such as transmission heterogeneity that can sustain malaria transmission indefinitely. While the ethics and safety of these methods are being debated, mathematical models offer an efficient way of predicting the behavior and estimating the efficacy of these interventions if deployed to specific regions facing challenges to reaching elimination. We have developed a detailed mathematical model of vector genetics where specific genomes code for physical attributes that influence transmission and are affected by the surrounding environment. This is the first model to incorporate an individual-based multi-locus genetic model into a detailed individual-based model of malaria transmission. This model opens the door to investigate a number of subtle but important questions such as the effects of small numbers of mosquitoes in a region sustaining malaria transmission during the low transmission season, and the success of gene drives in regions where extant vector control interventions could kill off gene drive mosquitoes before establishment. Here, we investigate the reduced efficacy of current vector control measures in the presence of insecticide resistance and evaluate the likelihood of achieving local malaria elimination using gene drive mosquitoes released into a high transmission setting alongside other vector control measures.

## Introduction

Malaria remains a deadly disease in a number of regions around the world but increased surveillance, improved access to care and vector control have put elimination in sight in a number of countries worldwide. In sub-Saharan Africa, where malaria is largely endemic [1], vector control is a cornerstone of control and elimination efforts, and insecticide based interventions such as insecticide treated nets (ITNs) and indoor residual spraying (IRS) are the most widely used vector control tools [2]. This has led to large decreases in malaria transmission in the region with ITNs being responsible for around 68% of averted cases [3].

However, the effectiveness of malaria control through insecticides is being threatened by growing insecticide resistance in a number of countries [2, 4, 5]. Of the 81 endemic countries surveyed between 2010 and 2018, 73 showed at least one major malaria species being resistant to at least one of four insecticide classes approved for malaria control [6]. Additionally, pyrethroids have thus far been the only approved class of insecticides for ITNs, and resistance to these insecticides is widespread, which severely compromises insecticide-based vector control [7]. To further complicate matters, mechanisms for resistance vary widely given the different target sites in the vector genome for different classes of insecticides and the differing decay rates of killing efficacy across insecticides. Point mutations result in reduced sensitivity of the mosquito nervous system to insecticides [8] while amplification or over-expression of certain genes that result in increased enzymatic metabolism of insecticides [9] is another form of resistance. Behavioral changes in a vector population within a few generations due to extreme stress exerted by the introduction of insecticides [10] has remained a more difficult form of resistance to identify and mitigate. There is a critical need to address these threats expeditiously to not lose ground in the fight against malaria.

The use of transgenic mosquitoes that carry gene drives has been proposed as another form of vector control [11]. The use of gene drives has been put forward as a means to address loss of intervention effectiveness in a region due to insecticide or drug resistance [12]. Furthermore, gene drives could be a potentially ideal modality to drastically reduce vectorial capacity [13] in high transmission settings where current vector control interventions under the most optimal conditions could fail to achieve elimination [14]. Gene drives could also address changing mosquito behavior such as increased outdoor biting due to increasing indoor vector control pressure [15]. A gene drive system based on preferential inheritance can result in an entire population acquiring an engineered genetic trait and a desired effect within a few generations [16]. Gene drive methods for the purposes of vector control broadly fall under two categories: first, modifying a population to make it refractory to malaria, a practice referred to as population replacement [17], and second, restricting the population of a specific subspecies, which is referred to as population suppression [18]. James et al. [19], and Hammond and Galizi [20] provide an overview of different gene drive strategies being currently considered or developed.

There are a number of challenges to be addressed before gene drives become an accepted tool for vector control. Besides technical challenges such as engineering genetically modified (GM) mosquitoes with reduced fitness costs and deploying gene drive mosquitoes in areas with existing vector control that could kill GM mosquitoes before establishment, addressing community concerns and communicating the ecological risks and epidemiological benefits of gene drives in a region are crucial to deploying gene drives to fight malaria [21].

Due to these challenges, mathematical models offer one of the best methods to evaluate the spread and impact of transgenic mosquitoes in a given setting. Examples of *in silico* models include population suppression by driving the Y chromosome or replacement using dual germline homing in different spatiotemporal settings [22, 23], optimal homing rates of multiplexed guide RNAs to reduce resistant alleles and increase the chances of population suppression or replacement [24, 25], and a reaction diffusion model to study fixation of deleterious gene drives through accidental release [26, 27]. These models include both agent-based approaches [22, 28], which are excellent for modeling small, isolated populations characteristic of a suppression drive, or continuous well-mixed populations [26, 29], which offer a rapid way of estimating long term effects of a gene drive campaign. Additionally, insecticide resistance has been modeled using either compartment models [30–32] or statistical approaches based on field data [33, 34]. However, given future release scenarios for gene drives into regions with existing vector control, insecticide resistance, distinct seasonalities, and specific physical barriers that could result in complex situations where small numbers of mosquitoes could be responsible for a strategy succeeding or failing, an agent-based approach to modeling vector genetics within the context of a vector-borne disease that includes all of these features would be invaluable.

With the aim of addressing all of these requirements we have developed, and describe in this article, the key components of a new stochastic, agent-based vector genetics model that follows Mendelian inheritance rules influenced by mosquito life and feeding cycle dynamics in a spatiotemporal setting. An agent-based modeling approach involves simulating individual agents such as humans or mosquitoes that follow both intra-agent and inter-agent rules. Intra-agent rules could govern phenomena such as the development of intra-host immunity or the expression of phenotypic traits by an individual based on their genotype while rules governing interaction between individuals also determine population-wide phenomena such as the transmission of disease. Key details about the differences between an agent-based approach to modeling malaria and mosquito dynamics versus other modeling approaches such as compartmental models can be found in a number of articles [35–38]. To orient the reader towards the working of this model, we then present examples of how phenotypes and genotypes interact in the model, how gene drives establish themselves in a population, and how species introgression may be captured in the model. We then proceed to more complex examples where the vector genetics model is embedded into an agent-based model of malaria dynamics. We demonstrate the impact on mosquitoes interacting with vector control interventions that are deployed in a region with high malaria transmission. We also look at factors for success of population replacement gene drives that result in vectors becoming refractory to malaria with or without the presence of vector control in a given setting, and the role of transmission heterogeneity and vector migration on malaria elimination efforts. Finally, we present future use cases of the vector genetics model ranging from vector control deployment strategies that combat insecticide resistance to optimizing gene drive releases to achieve malaria elimination.

## Methods

### Vector genetics model

Mosquitoes in EMOD v2.20 [39] can be modeled as individuals or cohorts. Detailed descriptions of how mosquito dynamics and malaria transmission are modeled in EMOD can be found in previous work [36, 38]. Each individual or cohort has a 64-bit diploid genome that is a recombination of two haploid genomes or gametes inherited from each parent, respectively. This diploid genome can account for up to 10 different loci or genes with up to 8 different alleles per gene. Four bits are reserved to code for the presence of microbial interventions such as *Wolbachia* [40] or other biological insecticides such as Metarhizium [41] (supplementary figure S1). Microbial interventions are not explored in this article but the modeling of *Wolbachia* and cytoplasmic incompatibility within EMOD has been covered in previous work [38]. When mating occurs in the model, the male and female produce offspring with a combination of genes and alleles obtained from the gametes of the parents via Mendelian inheritance (Fig. 1). Mutations can occur during gametogenesis (supplementary figure S2), and phenotypic traits can be assigned to specific genotypes (Fig. 1). Broadly, the traits that can be modified currently could affect the ability of a vector to transmit malaria, generate progeny biased towards a specific gender, confer fitness costs such as reduced fecundity or increased mortality, and simulate partially or fully insecticide-resistant vectors. The model also captures genetic drift due to factors such as spatial bottlenecks, initial allele skewness and fitness costs associated with environmental factors. By defining species complexes, the model can be extended to simulate subspecies introgression as well.

**Fig 1.**
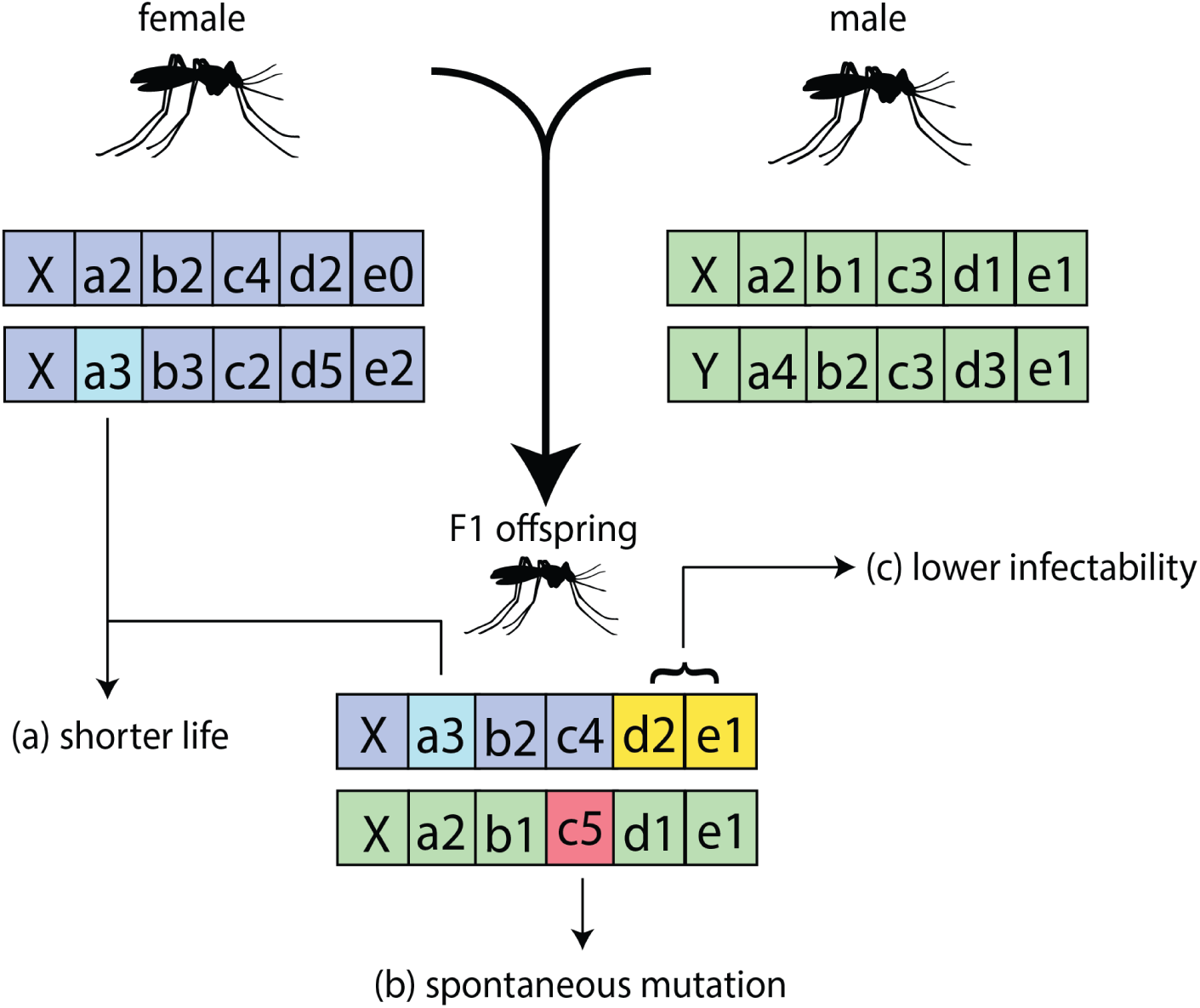
Representative genomes of cohorts or individual mosquitoes within EMOD. The model supports the inheritance of traits from parents (a), random mutations of alleles (b), and definitions of phenotypic traits associated with a combination of genes or alleles, that are expressed only when those combinations are present (c). Inheritance of genes is modeled as a Mendelian process, and combinations of alleles can be mapped to combinations of traits via a many-to-many mapping.

### Gene drives within EMOD

Various gene drive strategies can also be simulated using this model such as classic endonuclease drives where driver and effector genes are driven as one construct [42], integral drives with independent autonomous driver and non-autonomous effector genes [43], as well as daisy-chain drives with serially dependent but unlinked drive elements [44]. In the model, gene drive dynamics are based on endonuclease drives, which work by cutting a chromosome at a specific location that does not include the drive prompting the cell to repair the cut with a copy of the drive. Godfray et al. [23] provide a detailed explanation of the mechanism of endonuclease gene drives. Cleaving at the target site and copying of the desired allele occurs during gametogenesis (Fig. 2). For Mendelian inheritance rules, see supplementary figure S2. This allows for simulated mosquitoes to carry the drive in a heterozygous or homozygous configuration to facilitate modeling fully or partially recessive traits associated with driven alleles, as used in either population suppression or replacement contexts.

**Fig 2.**
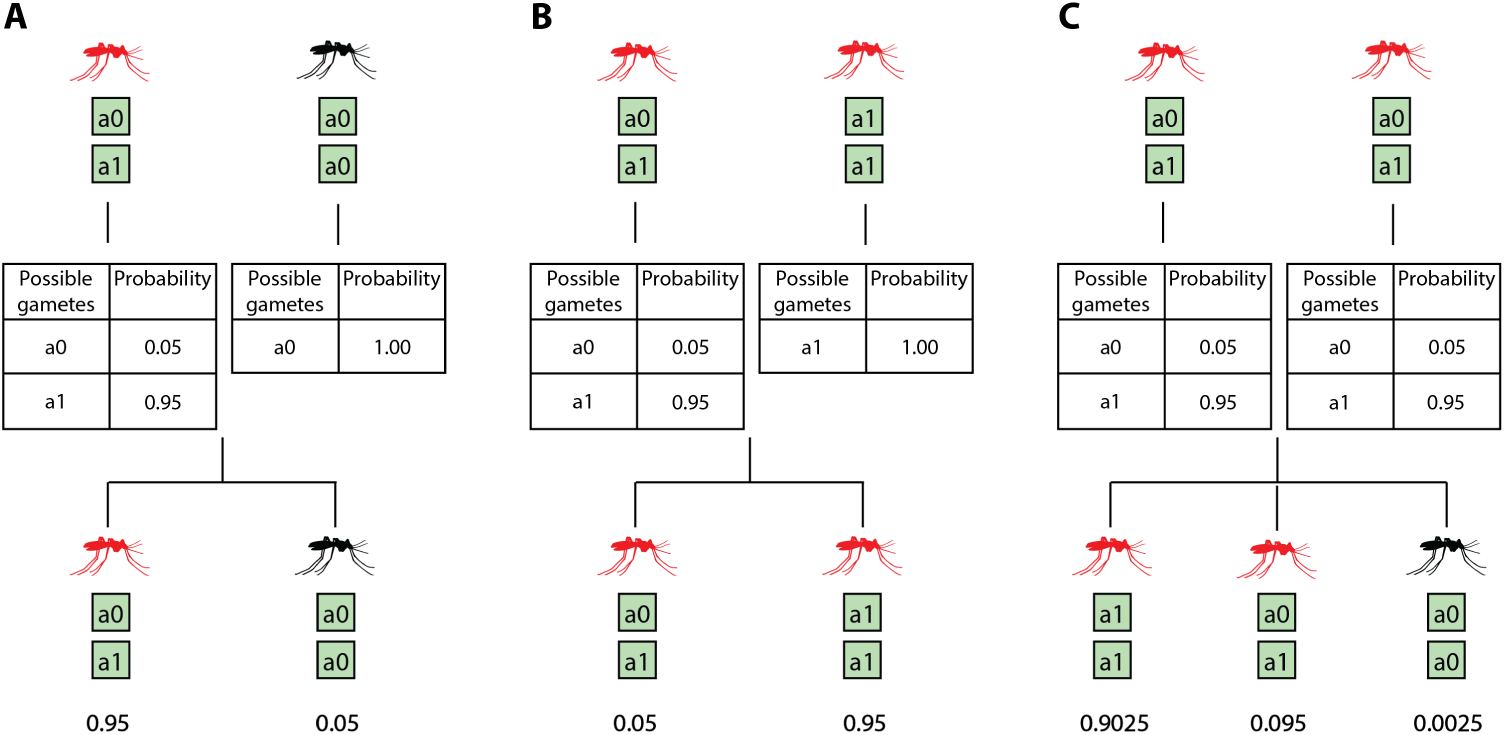
Inheritance rules for gene drives. In this example, ‘a1’ is the driven allele and ‘a0’ is the wild-type allele. Red mosquitoes represent mosquitoes with the driven allele ‘a1’ either in the homozygous or heterozygous configuration. Black mosquitoes represent wild-type mosquitoes, which are homozygous in ‘a0’. The drive successfully cleaves the target site 95% of the time. In the event of drive failure, the wild-type allele remains. This is akin to modeling drive failure in terms of the driven allele failing to home in to the target site. This does not include target site resistance through natural mutation or non-homologous end joining (NHEJ). No other mutations are considered. Three mating scenarios are presented: (A) When a drive carrying heterozygous mosquito mates with a homozygous wild-type mosquito, (B) a heterozygous-with-drive mosquito mates with a homozygous-with-drive mosquito, and (C) two heterozygous-with-drive mosquitoes mating.

### Insecticide resistance within EMOD

Insecticide resistance is modeled as a phenotypic trait associated with the expression of specific alleles or allele combinations in an individual vector or a cohort of vectors. Resistant alleles are defined for each insecticide at the start of the simulation. The expression of resistant alleles in a vector could code for changes in either killing efficacy or the efficiency of repellence of insecticides applied to various interventions such as insecticide treated nets (ITNs), indoor residual spraying (IRS), attractive targeted sugar baits (ATSBs) and space spraying. Additionally, vector host seeking behavior can also be modified to model behavioral resistance. Fitness costs affecting the lifespan of vectors or their fecundity can also be imposed and associated with combination of genes and alleles.

### Simulation framework

All simulations were carried out with EMOD v2.20 [39], an agent-based mechanistic model of *Plasmodium falciparum* malaria transmission with vector life cycle [38], and parasite and immune dynamics calibrated to within-host asexual and sexual stages of the parasite [45]. The vector life cycle in the model consists of four stages : eggs, larvae, immature adults and host seeking adults. The larval habitat available at a given time dictates the total number of larvae that can be accommodated per habitat type at that time, which in turn governs the number of vectors emerging at the end of the life cycle.

For simulations modeled as a single location, a single peak seasonality characteristic of the Sahel [45–47] is used to model the amount of vector habitat available at a given time (Fig. 3A). Malaria transmission is modeled in a population of approximately 1000 individuals with birth and death rates observed in the Sahel, and an effective annual entomological innoculation rate (EIR) of 120 absent any interventions [45]. For multi-location spatial simulations, population data was obtained using the High Resolution Settlement Layer (HRSL) generated by the Facebook Connectivity Lab and Center for International Earth Science Information Network (CIESIN) at Columbia University [48]. An area covering 300 square kilometers in rural Burkina Faso was resolved into one square kilometer grid cells, and only grid cells with more than 5 people were assumed to be inhabited, which resulted in a total of 150 populated grid cells and total of 4000 individuals (Fig. 3B).

**Fig 3.**
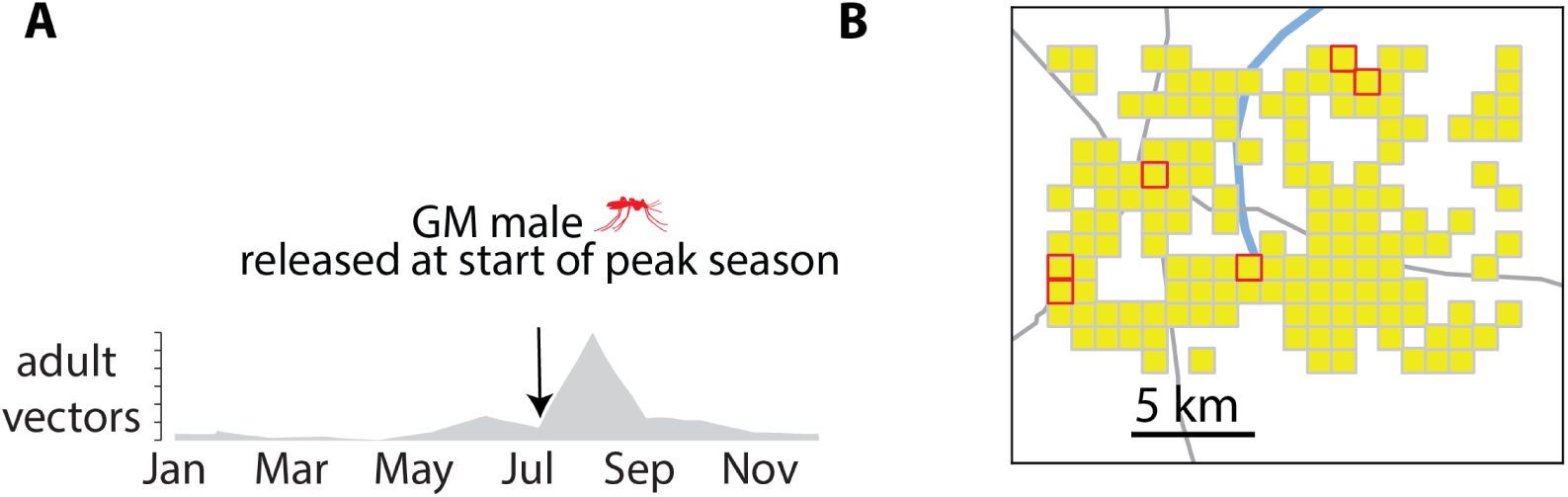
Seasonality and spatial simulation setup using EMOD. (A) Single peak seasonality profile characteristic of a Sahelian seasonal and transmission setting. All simulations presented here with a prevalence endpoint measure use this seasonality profile resulting in an annual EIR of around 125 infectious bites per person. (B) Spatial grid used for spatiotemporal gene drive simulations. Six nodes with the largest human population are selected as release sites for genetically modified mosquitoes carrying drives and are marked in red.

Human individuals in these spatial simulations are assigned a daily probability to take overnight trips to other grid cells according to a distance decay relationship [49]. The distance decay is calibrated to human movement on scales of one to tens of kilometres observed in geotagged campaign data [50] leading to an average of 5 overnight trips per person per year. There is no disease importation from outside the modelled area. vector carrying capacity in each node is scaled to human population such that humans have the same probability of being bitten across all nodes. Vector migration also follows the same distance decay model with migration rates scaled by the ratio of total vector to total human population, and with preferential migration to nodes with higher larval habitat. There is no human or vector migration into or out of the modeled area.

Fitness costs can be modeled using the genotype to phenotype mapping feature, as demonstrated for the simple gene drive example in figure 6. However, in every other gene drive example, we consider all genotypes to be equally fit To show the maximum impact achievable on malaria transmission by gene drives.

**Fig 4.**
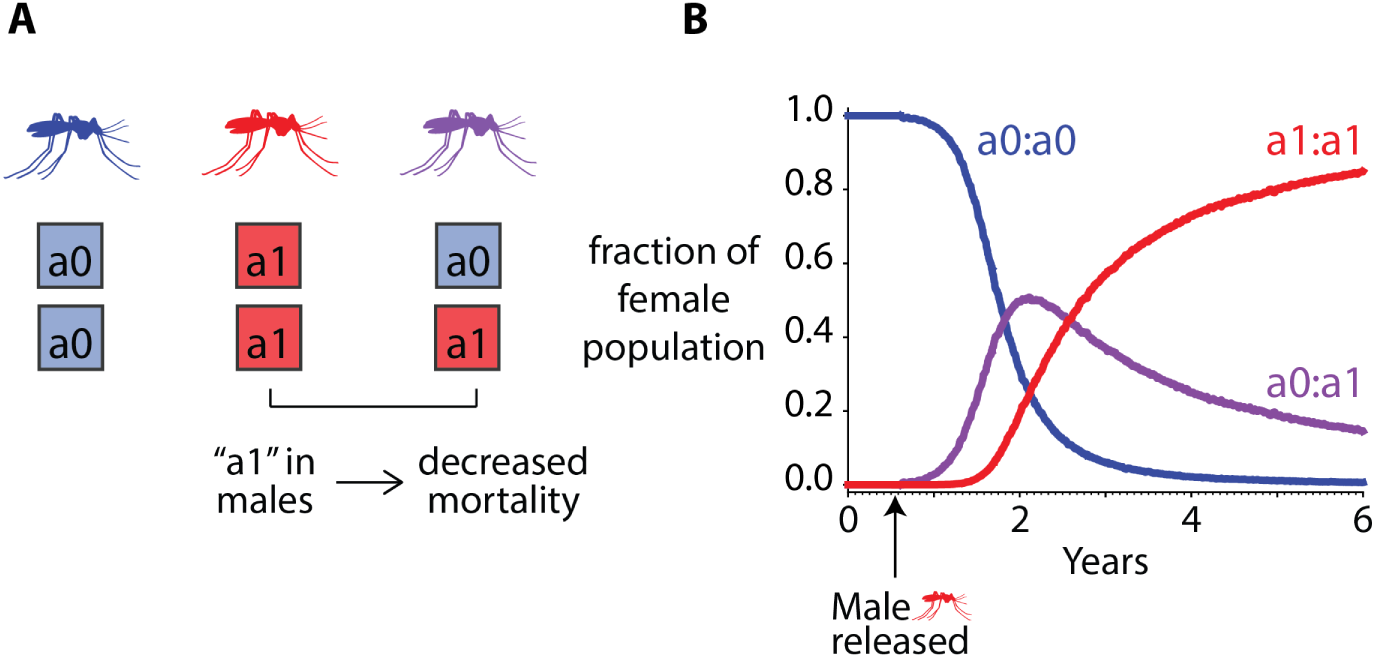
Example of how traits and alleles interact in the vector genetics model. There is no seasonal variation and no spatial component. (A) Genomes of mosquitoes in the model. Mosquitoes homozygous in ‘a0’ are the wild-type mosquitoes. Male mosquitoes carrying the ‘a1’ allele in a homozygous or heterozygous configuration have decreased mortality, which is thus a dominant trait, and is modeled as a halving of their probability of dying. (B) Distributions of genomes in the population over time average over 50 stochastic realizations. Male mosquitoes homozygous in the ‘a1’ allele are released mid-year during the first year of the simulation.

**Fig 5.**
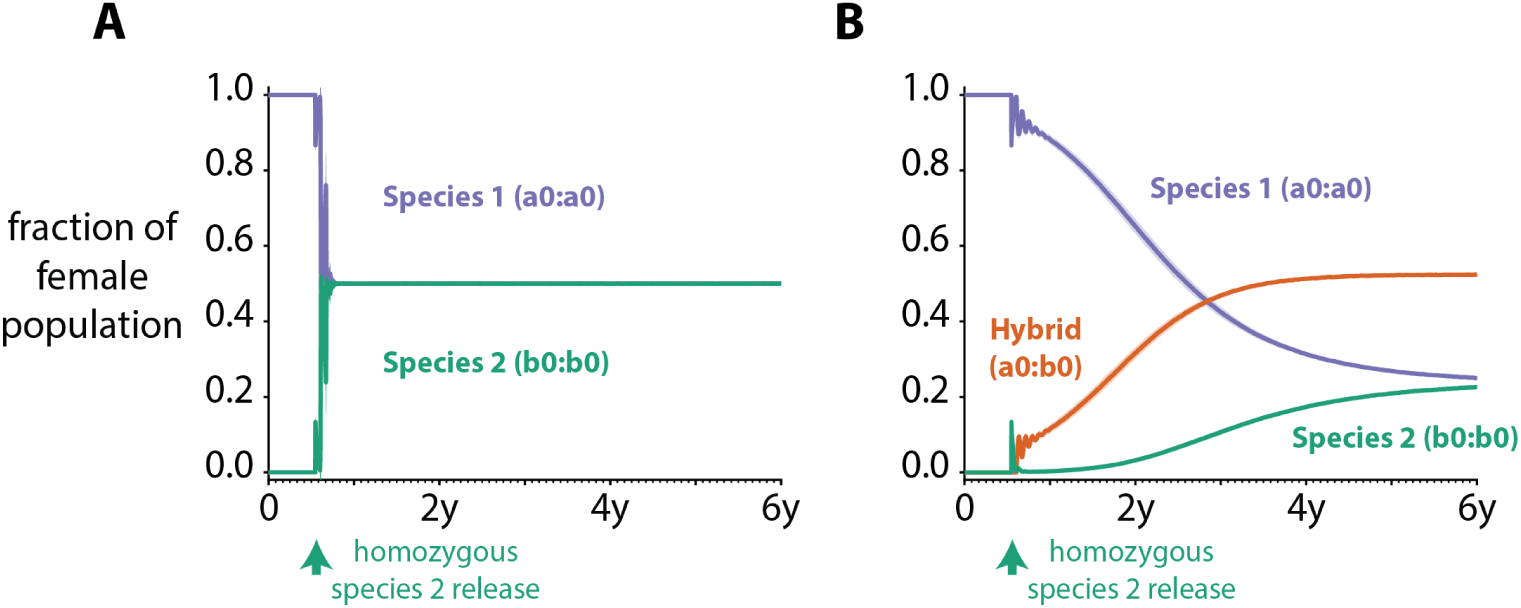
Example of species introgression in the vector genetics model. The mean (solid line) and one standard deviation (shaded area) of genomes in the population over time of 50 stochastic realizations when 10000 male and female mosquitoes of species 2 that are homozygous in ‘b0’ are introduced into a population containing only species 1 mosquitoes homozygous in ‘a0’ 200 days after simulation starts. (A) When no introgression occurs, and each species has equal habitat to procreate. (B) Hybrid mosquitoes heterozygous in ‘a0’ and ‘b0’ have a 10% lower mortality rate than species 1 or species 2 mosquitoes.

**Fig 6.**
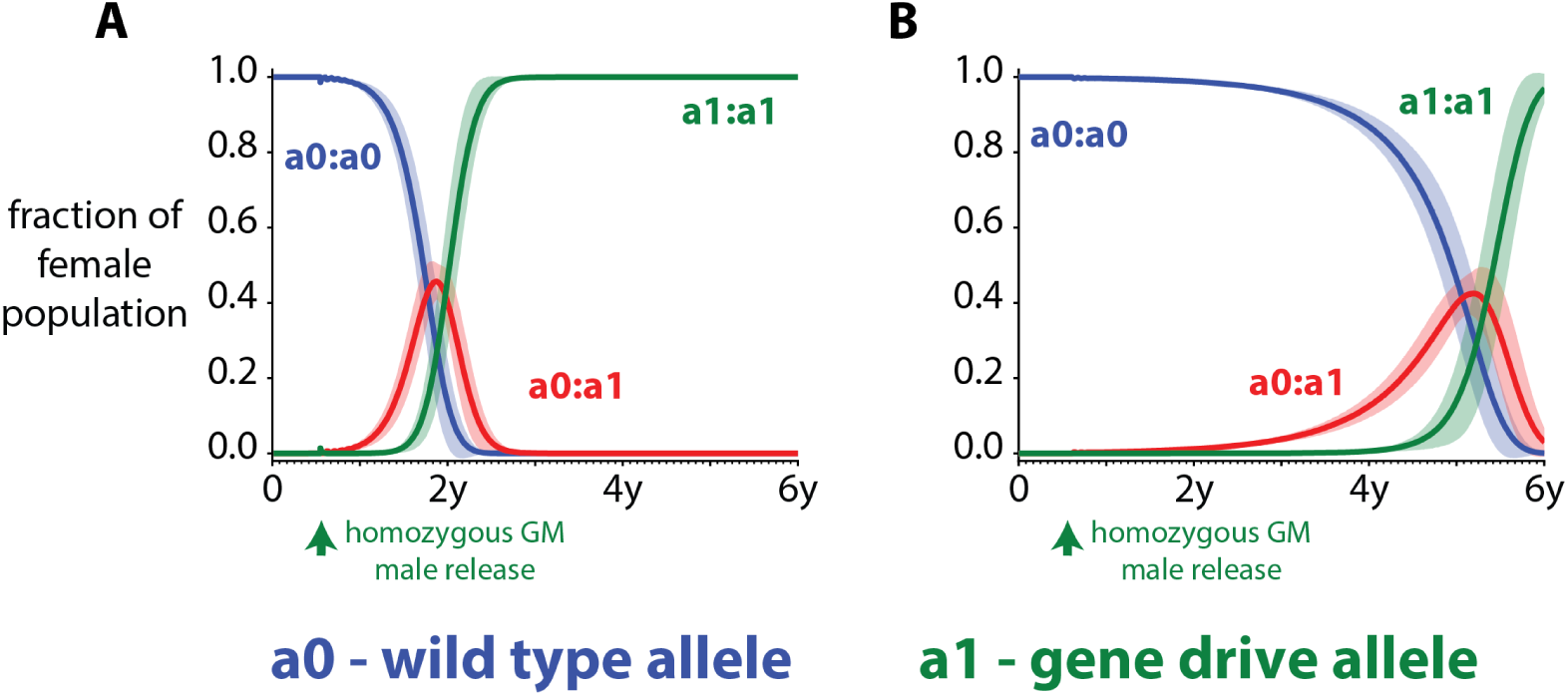
Example of how gene drives behave in the vector genetics model. The mean (solid line) and one standard deviation (shaded area) of genomes in the population over time of 50 stochastic realizations when male mosquitoes with drive are released into a wild-type population. There is no seasonal variation and no spatial component. Mosquitoes homozygous in ‘a0’ are the wild-type mosquitoes. Male mosquitoes carrying the ‘a1’ allele in a homozygous configuration are released 200 days after simulation starts. The ‘a1’ allele is a driven allele and has a homing rate of 50% to the ‘a0’ site. (A) There are no fitness costs associated with drive mosquitoes. (B) Mosquitoes with drive in either a homozygous or heterozygous configuration have a 10% higher mortality rate than wild-type mosquitoes.

### Interventions

Treatment with artemether–lumefantrine (AL) is available for symptomatic cases in all simulation scenarios. 80% of severe malaria cases are assumed to seek treatment, and treatment is sought within 2 days of symptoms occurring. We assume only 50% of clinical cases seek treatment, which happens within 3 days of symptom onset [51].

Intervention scenarios are simulated for six years. ITN distributions occur every three years per WHO recommendations [52] on July 1 just as the peak season starts to pick up. ITN retention is modeled as decaying with exponential rate of two years [3]. In the insecticide resistance scenario, ITN killing efficacy starts at 80% with an exponential decay rate of four years per WHO guidelines for classifying a vector population as susceptible. In the gene drive scenarios, ITN killing starts at 60% to account for efficacy loss due to insecticide resistance [53]. A fixed killing rate is chosen to keep these scenarios focused on the gene drive aspect of the vector genetics model. ITN blocking starts at 90% with an exponential decay rate of two years to model the physical integrity of nets for all scenarios [53].

In the gene drive scenarios, mosquitoes carrying a population replacement drive are released on July 1 just as the peak season begins to pick up to give the drive maximum chance of spreading. In the single location simulations, the likelihood of elimination is evaluated for different levels of transmission efficacy from mosquitoes to humans, and drive cleave-and-copy success probability. The efficiency of the drive, that is the copying over of the drive allele at the target locus, is varied from 50% to 100%. The probability of infected vectors carrying the drive transmitting malaria to humans is varied from 0 to 50% compared to a vector without drive. For the spatial simulation, the likelihood of drive copying over is maintained at 100% while the probability of transmitting malaria to a human when the vector carries the drive is reduced to 30% compared to a vector without drive. These parameter values were chosen in the spatial simulation to highlight any differences between the multi-node and single node simulations that may arise from the migration of mosquitoes.

Given the absence of importation or migration outside the simulated area, elimination is defined as zero infected individuals at the end of the sixth year. Each scenario is run for 50 stochastic realizations.

## Results

In each of the simulation scenarios, values for the parameters within the model are chosen within physiologically plausible limits. Additionally, these values were chosen to demonstrate the capabilities of the model and reflect malaria transmission in a Sahelian setting. Calibration of the model to entomology, genetics, and epidemiological data representative of a specific setting are required before simulating future scenarios and outcomes that are predictive of the same setting.

### The vector genetics model can capture the effects of fitness costs and benefits of specific phenotypes on an entire population as well as the inheritance of gene drives in the model

To demonstrate how different mosquito phenotypes interact with each other in the model, we released 1000 male mosquitoes homozygous with a mutated allele, which reduces the mortality of male mosquitoes by half (Fig. 4A), into a wild-type population of 100000 adult male and female mosquitoes each. The reduced mortality is modeled as a dominant trait conferring the same phenotypic characteristics to the male mosquito irrespective of its zygosity with respect to the mutated allele. For this single location simulation, no seasonality was imposed to isolate conferred physical traits as the cause for propagation of different alleles in the population. The modified males were released six months after the start of the simulation (Fig. 4B). We have chosen not to include mutations in any of the genotypes in this simulation to focus solely on the effects of fitness benefits.

Just under a year after the start of the simulation, heterozygous females start to emerge. A year after the release of the male mosquitoes carrying the mutated allele, homozygous in ‘a1’ females start to emerge. As the simulation progresses, the mutated ‘a1’ allele dominates the population while the ‘a0’ allele-carrying females die out. Males carrying the ‘a1’ allele either in a homozygous or heterozygous configuration live longer and have a greater chance of passing on their genes to offspring. This results in the ‘a1’ allele being propagated to almost the entire population five years after release.

Introgression of genes from different species may also be accounted for in this model. If genes from one species do not introgress into the genome of another, and each species has separate but equal access to breeding sites, we see a quick redistribution of genome frequencies to reflect a 50-50 split of vectors in a site (Fig. 5A) when a new species is introduced. If, however, genes from the newly introduced species may introgress with the existing species in the site, we see the emergence of a new hybrid population. Adaptive introgression has been observed in many species including mosquitoes as a result of factors such as insecticide pressure or environmental changes due to, for example, anthropogenic climate change, and could confer phenotypically advantageous characteristics to the emergent hybrid population (Norris et al., 2015). In figure 6B, the hybrid population has a 10% lower mortality rate than both the native and introduced species, and becomes the dominant mosquito in the region just over 2 years after the introduction of Species 2. In addition to the fitness benefit of the hybrid mosquito, all three subspecies of mosquitoes – Species 1, Species 2, and hybrid – compete with each other for existing larval habitat, which dictates the final establishment rates of each genotype in the simulation.

As an example of how gene drives work in the model, we released 1000 male mosquitoes carrying a gene drive allele ‘a1’ in a heterozygous configuration into a population of 100000 male and female mosquitoes carrying the wild-type ‘a0’ allele in a homozygous configuration. The ‘a1’ allele cleaves and copies the ‘a0’ allele at the target site of the second chromosome with a success rate of 50%. This is representative of drive failing to home in on the target site 50% of the time. There are no mutations modeled. Males carrying the gene drive allele are released 200 days after the start of the simulation. Again, no seasonality was imposed in this single location simulation (Fig. 6).

Heterozygous females start to emerge almost immediately after release and just as the homozygous in ‘a1’ females start to form a large fraction of the population about a year after the release, there is a precipitous drop in the wild-type allele, ‘a0’, in the population (Fig. 6A). A homing rate of 50% results in the ‘a1’ fixating in the population around 2 years after the initial release of GM mosquitoes carrying gene drives. In the absence of fitness costs, vector migration, and seasonality, the ‘a1’ allele will always fixate but the rate at which it achieves fixation will be dependent on the homing rate. Fitness costs, on the other hand, delay establishment of the drive. For example, a 10% increase in mortality due to the drive could result in fixation occurring almost 6 years after release instead of 2 years (Fig. 6B).

### Insecticide resistance could lead to malaria transmission rapidly becoming refractory to vector control that involves repeated deployment of the same insecticide

Vector control tools such as ITNs and IRS have been deployed extensively across the Sahel over the past two decades. In a high transmission setting with seasonality modeled after the single peak wet season (Fig. 3A) the start of the peak season is the most optimal time to deploy ITNs [45]. In the modeled scenario presented here, vectors heterozygous with a resistant allele constitute around 3% of the total vector population while vectors homozygous with a resistant allele constitute less than 0.1% of the total population at the start of the simulation. The heterozygous vectors are modeled as being partially resistant to insecticides with killing efficacy reduced to 10% when they interact with nets while vectors homozygous with the resistant allele are fully resistant with only 5% efficacy in killing. Please refer to the interventions subsection of the methods section for intervention efficacy with respect to wild-type vectors. When nets are deployed at the end of June, there is rapid selection of the resistant allele (Fig. 7B). However, the effect of increasing the resistant population on malaria transmission is only observed after the second deployment of ITNs treated with the same insecticide three years later (Fig. 7).

**Fig 7.**
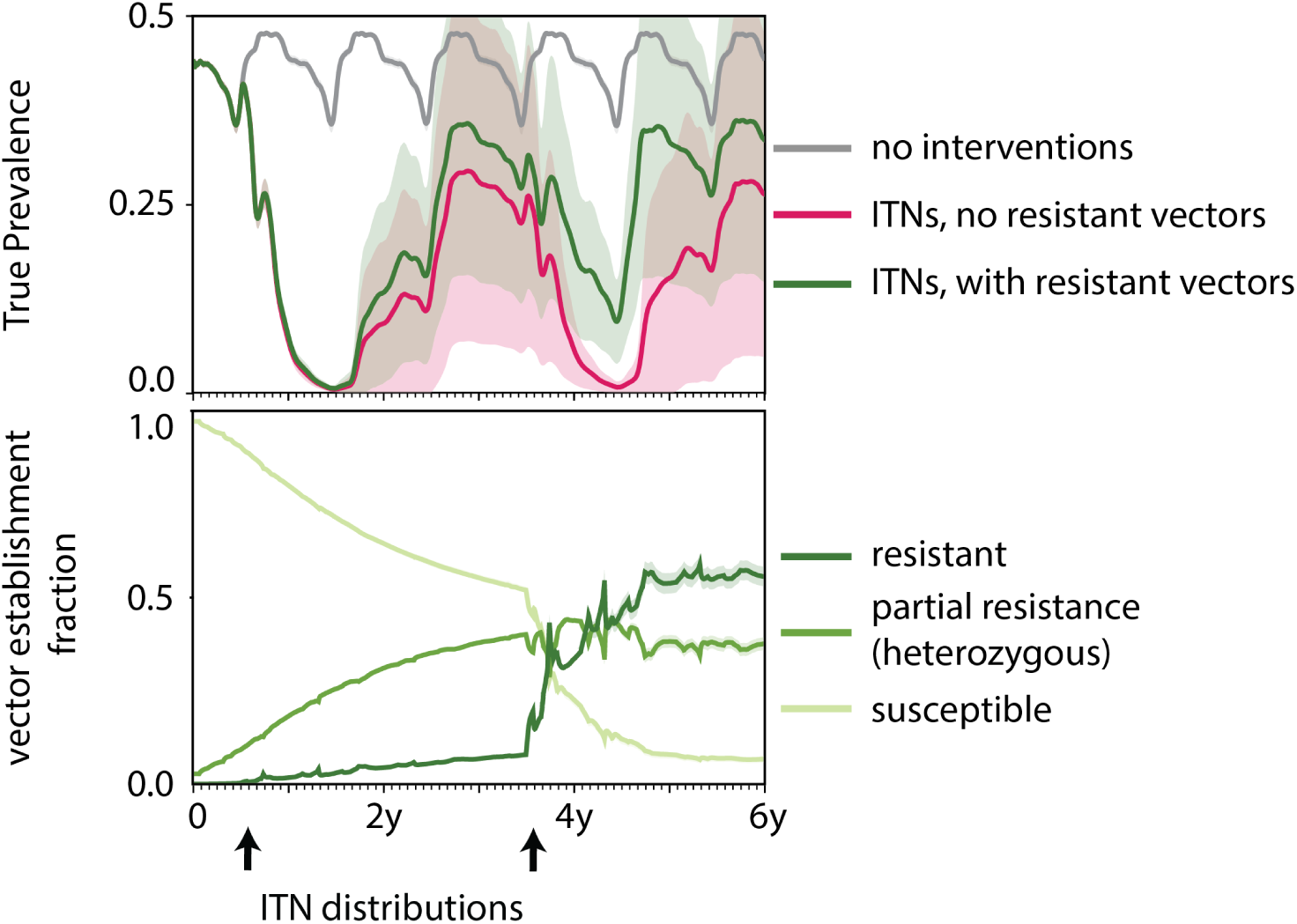
Tracking the effect of insecticide resistance on prevalence in a Sahelian setting. RDT prevalence (top row) across a period of six years averaged over 50 stochastic realizations for three scenarios – when no interventions are deployed, when ITNs are deployed at 60% coverage absent any resistance, and with the presence of resistance. Establishment rates of susceptible and resistant genomes in the scenario with resistance (bottom row). The shaded area around the mean represents one standard deviation calculated across the 50 stochastic realizations. ITNs are distributed every three years at the beginning of the peak season.

When no interventions are deployed RDT prevalence oscillates between 15% in the low season to a high of around 45% in the peak season. However, when ITN’s are deployed with no resistance, peak RDT prevalence is around 10% in the middle of the third year and drops to levels under 2% after the second deployment of nets. As usage wanes towards the end of year 6, RDT prevalence peaks at 5%. The total number of clinical cases, however, has dropped by 85% over the course of six years resulting in an annual EIR of 9 infectious bites per person. In the case with resistance, RDT prevalence mimics the scenario with no resistance until the fourth year. Selection of resistant vectors increases the resistant proportion of the population (supplementary figure S3) leading to higher sustained prevalence rates similar to rates seen after the first deployment of nets with a peak of 10%. The total number of clinical cases averted over the six years drops to 80%. The annual EIR climbs to a mean of 30 infectious bites per person over the course of the final year.

### Traditional vector control strategies increase the chance of malaria elimination via genetically modified vectors refractory to malaria in high transmission settings

In highly seasonal settings, releasing mosquitoes that carry gene drives at the start of the peak season maximizes the chances of spreading introduced genes through the entire population [54]. Here, vectors carrying a gene drive that renders them refractory to malaria are released at the end of June in a setting with seasonality characteristic of the Sahel (Fig. 3A). The spread and effect of malaria refractory gene drives in inhibiting malaria transmission is explored in three scenarios: one with no other forms of vector control, and two scenarios with ITNs distributed at the start of the wet season every three years at 60% and 80% coverage, respectively. This is a single location simulation and for each scenario the probability of elimination six years after the start of the simulation is calculated for a range of transmission blocking efficiencies and successful drive copy rates (Fig. 8).

**Fig 8.**
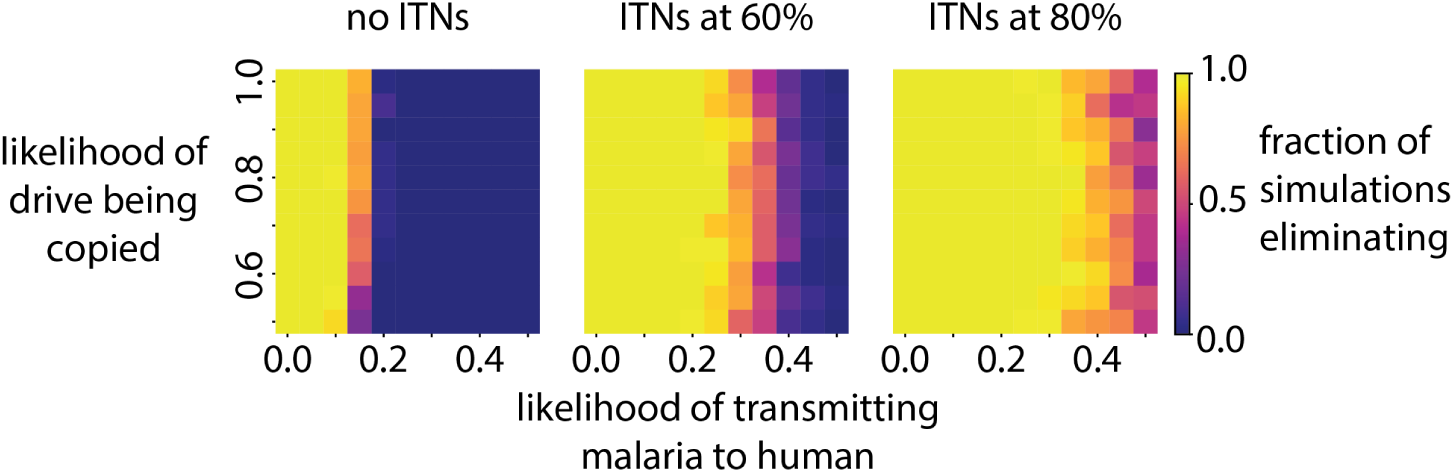
Likelihood of elimination in a Sahelian setting using gene drives that reduce the probability of transmission of parasites from mosquitoes to humans. Three scenarios are presented – one without ITNs, and ITNs distributed every three years over a six year period at 60% and 80% coverage, respectively. The fraction of simulations eliminating is evaluated over 50 stochastic realizations for a given value pair of probability of transmission from mosquito to human and likelihood of successful gene drive of the gene responsible for reduced transmission.

In the use case of a population replacement gene drive, the likelihood of the driver successfully cleaving the target site and copying over the desired gene has minimal impact in all three scenarios. However, it does have an impact on how quickly the drive establishes (supplementary figure S4). This in turn has an effect on how quickly transmission is reduced. However, the degree to which the desired gene blocks transmission of the malaria parasite from mosquito to human is a stronger predictor of elimination. When no ITNs are present, and when the likelihood of infected vectors transmitting malaria to humans is greater than 20%, malaria persists in the region. However, with the addition of ITNs at 60% coverage, a vector to human transmission efficiency of 35% still results in 50% of the simulations eliminating. At 80% ITN coverage and only 50% chance of a bite being infectious, there is a 60% chance of elimination in the region. In the last two scenarios, the inclusion of ITNs increases the chances of elimination despite ITNs killing off vectors carrying the drive as well. The drive spreads through the population because male mosquitoes are unaffected by ITNs and reseed the vector population with the drive until the drive establishes in the entire population.

### Spatial connectivity and interaction with traditional vector control are critical factors in determining gene drive success

Vector migration plays an important role in the spread and establishment of a gene in a regional vector population as migration determines the rate of gene flow between subpopulations that are spatially segregated. To explore the effects of a region’s connectedness and the migration of vectors between habitat and meals, the Sahelian seasonality (Fig. 3A) is imposed on a representative region of settlements in the Sahel (Fig. 3B). Release sites for mosquitoes carrying drive are marked in red, and 100 genetically modified mosquitoes are released from each of the sites at the end of June as the peak season begins to pick up. The effects of a vector control intervention such as ITNs when combined with a gene drive release are explored (Fig. 9).

**Fig 9.**
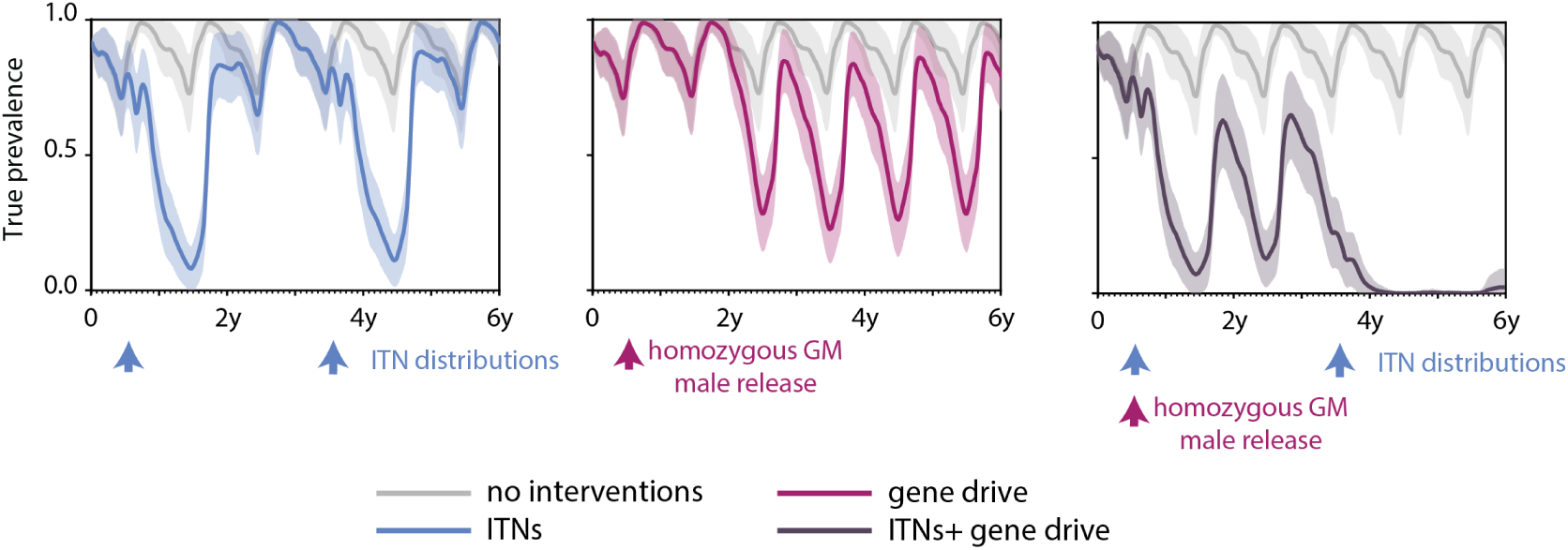
Evolution of true prevalence over time when different intervention packages are deployed in a multi location Sahelian setting. Results are from a spatial simulation with 150 one square kilometer nodes with varying population and larval habitat sizes spread across 300 square kilometers. Four different scenarios are explored – absent any vector control interventions, release of genetically modified mosquitoes carrying a drive that reduces the probability of parasite transmission from mosquitoes to humans, deploying ITNs every three years just as the peak season begins to pick up, and a combination of gene drive and ITNs. The average of 50 stochastic realizations of each scenario is represented by solid lines while the shaded area represents one standard deviation.

For gene drive releases in the multi-location simulations, infectious vectors carrying the drive are modeled to have a 70% drop in their efficiency to transmit malaria to a human. The likelihood of drive copying over is maintained at 100% to simulate an optimistic scenario for gene drive establishment in this setting. Vector migration is modeled as described in section Simulation framework. When there are no interventions, the true prevalence in the region oscillates between around 65% in the dry season to over 95% at the peak of the wet season. With the introduction of ITNs at 80% coverage, large drops in prevalence are observed immediately after deployment but waning usage of the nets over time coupled with decreasing net integrity and insecticide effectiveness over time leads to prevalence reverting to levels without vector control a year and a half following deployment. This is seen after both ITN distribution events that are three years apart.

In the scenario where there is a single release of mosquitoes carrying drives at the end of June in the first year of the simulation, there is a large drop in prevalence approximately 2 years after release. This is because vectors carrying the drive take time to migrate away from the release sites and establish in other areas before driving down transmission. However, after establishment, transmission persists with a maximum prevalence of around 90% and a minimum prevalence of around 25% over a given year. This is in line with single location simulations of gene drive releases (Fig. 8). Additionally, the release sites have been held constant at six for this scenario. In this setting, the number of release sites only impacted the speed of establishment and not the probability a gene drive would establish. Six sites with the largest human population were chosen to keep release numbers under 1% of the total vector population in the simulated region.

However, when gene drives and ITNs are combined the probability of elimination decreases in the multi-location simulation compared to the single location ones. In the multi-location scenarios, gene drive mosquito release and the first round of net distribution are conducted concurrently at the end of June of the first year of simulation with an additional round of net distribution after three years. Again, gene drives take two years to establish and have an impact on transmission but ITN usage in the meantime drives the prevalence lower in year 3 of the simulation than in the case with only ITNs. As the impact of gene drives begins to grow after year 3, the prevalence drops to almost undetectable levels in combination with the second ITN distribution event. However, as ITN usage wanes, the prevalence begins to pick up again towards the end of year 5. Now only 80% of simulations eliminate as opposed to 100% elimination seen in the single location. Partial suppression of transmission due to drives and uneven migration leads to pockets of lower establishment (supplementary figure S5). The decreases in establishment rates are especially distinct around the start of the wet season when wild-type mosquitoes could have survived in greater numbers than GM mosquitoes because of demographic stochasticity, which small populations are vulnerable to, leading to a resurgence of wild-type mosquitoes in the node. This combined with dropping ITN usage lead to regions of high prevalence in some simulations (supplementary figure S5). This in turn leads to prevalence increasing if transmission is not eliminated by the time net usage starts to decrease.

### Computational resources and simulation times

All simulations were carried out using the Institute for Disease Modeling’s in-house high performance computing job management platform with access to up to 768 logical cores with a clock speed of 3.30GHz and 8GB of memory per core. In the single node simulations, approximately 200,000 male and female mosquitoes were simulated for 6 years with an average run time of 4 seconds per simulation. In the single node insecticide resistance and gene drive scenarios, 1.5 million male and female mosquitoes with a human population of 1000 individuals complete with malaria control interventions and individual immune dynamics were simulated for 6 years with an average simulation time of 8 minutes per simulation. In the spatial simulations, we simulate approximately 5000 people and 6 million male and female mosquitoes spread over 150 nodes using 5 compute cores with an average simulation time of 14 minutes per simulation. The spatial simulations include migration of humans and mosquitoes between nodes as well as malaria control interventions such as ITNs and health-seeking by infected individuals.

## Discussion

Insecticide resistance threatens to drastically undermine malaria control and elimination efforts globally. New tools are required to understand the spread of insecticide resistance, and how new technologies such as gene drives, which aim to overcome the challenges posed by insecticide resistance, should be optimally deployed in the field. Here, we present a mathematical model that captures gene flow, the impact of gene drives, and the evolution of resistance in vector populations under insecticide pressure in an agent-based spatiotemporal framework of malaria transmission.

The results from embedding this agent-based model of vector genetics within an agent-based model of malaria transmission dynamics show that even a small number of insecticide-resistant vectors can start to dominate a local population facing repeated exposure to the same insecticides. This could lead to a rapid rise in the number of clinical cases of malaria. Additionally, in regions with highly seasonal and heterogeneous transmission, existing vector control methods such as ITNs can work in concert with gene drives that seek to replace the wild population with mosquitoes that have a reduced ability to transmit the disease to bring even high transmission settings close to elimination in a few years.

The scope of this study is limited to presenting a modeling framework. To this end, we use parameter values that are within physiologically plausible limits and the examples presented here reflect the dynamics of vector genetics and malaria in a high transmission setting. Careful calibration to field data from a specific location is necessary to answer questions relevant to that setting. However, irrespective of differences between transmission settings, there are a number of unanswered questions with respect to mosquito gene flow, insecticide resistance and gene drives that mathematical modeling and this model in particular could aid in answering across a range of transmission intensities. For example, the model presented here could be leveraged to understand the rise of insecticide resistance given different vector control strategies in a region. For an assortment of insecticides that can be delivered via different modalities such as IRS and ITNs, mathematical models of resistance such as the one presented here could be used to calculate optimal timing and spatial deployment of these interventions as well as develop insecticide cycling strategies [55] that could mitigate the spread of resistant vectors. Furthermore, the impact of vector control strategies that employ two or more insecticides [56] deployed concurrently amidst resistance in a given region can also be assessed.

Another topic of future research involves behavioral resistance. Here, we have focused on a loss of killing efficacy for insecticides when resistance is present but changes in vector behavior due to resistance could lead to a further increase in transmission. For example, resistant vectors that are averse to insecticides may avoid landing on nets or entering houses treated with IRS [57] and shift transmission modes by preferentially seeking more outdoor feeds. While we have not explored these questions in detail in this work, the scenarios presented here serve as an example of how the model can be adapted to address scenarios arising from more complex resistant vector behavior.

The model is limited to 8 alleles and 10 loci to make the model computationally efficient by limiting the amount of memory allotted to each genome to 64 bits, which is a common width for registers in a CPU. Additionally, a 10 loci genome is sufficient to model a number of complicated scenarios ranging from single gene mutations in *kdr* resistant mosquitoes [58] to complicated multi driver effector gene drives that could be developed using emerging constructs [43]. There are, of course, limitations to this model such as the absence of a framework to account for genetic linkage of resistant alleles with alleles coding for other phenotypic characteristics, which could impact how genomes are selected for under insecticide pressure especially in the case of polygenic resistance. However, the modular framework of this model is extensible to include these characteristics should the need arise.

As the debate continues on if and how gene drives should be released, mathematical models could prove invaluable in narrowing down questions and concerns of regulators and stakeholders to aid in making informed decisions. The optimal size and timing of release that result in drive fixation given a deliberate or accidental release in a region, whether replacement, suppression or a combination of the two strategies is best suited to a region, and the effects of ongoing vector control on gene drives and their combined effect on malaria transmission are just a few examples of more specific questions this model can be leveraged to simulate within the framework of malaria transmission in a region.

In the past, we have used EMOD to answer complex question about the impacts of seasonality, transmission heterogeneity, and various intervention measures on malaria transmission [45, 49, 59], but we now have the ability to do so through the lens of vector genetics, and specifically, the outcomes of deploying complex intervention strategies such as gene drives. The power of this model thus lies in its ability to provide specific answers in detailed but limited geographies of the order of several hundred to a few thousand square kilometers encompassing populations of hundreds of thousands of people and several million mosquitoes. This limit is imposed to optimize computational efficiency while also providing actionable results to key stakeholders ranging from malaria research groups to national malaria control programs.

While we aim to use this model to investigate important questions related to vector genetics, we also aim to continually improve the model to accurately capture how resistance spreads or gene drives establish themselves in a region. For example, we simulate mosquito migration at shorter scales compared to phenomena such as long range migration aided by wind [46]. Additionally, we have limited migration to within the simulated region while there could be mosquitoes entering or escaping the region with dire consequences. Gene drive mosquitoes escaping the simulated or control region and establishing elsewhere is a genuine cause for concern [19]. These migration events could also impact transmission. For example, importation of wild-type mosquitoes into the study area could reseed transmission year after year. Or preferential migration to some nodes during certain times of the year could lead to other regions being primed for colonization by non-gene drive mosquitoes. Some of these questions have been tackled by other research. For example, the spread of gene drives and vector genetics over larger spatial scales has been modeled by North et al. [60]. However, we believe the full potential of the model presented here is realized when it is deployed to answer questions about malaria transmission and vector genetics at the sub-national scale; to the best of our knowledge, this is the first model to do so holistically. Additionally, while we have eschewed migration of vectors into and out of the simulated area to keep the examples simple, these scenarios may still be investigated within the current framework. For example, migration of vectors into the modeled area can be achieved in two ways: 1) as a release of mosquitoes as shown in the spatial example or 2) as migration from a high vector density node that is separate from the simulated area and reseeds mosquito populations in the simulated area seasonally or perennially. Similarly, migration of vectors out of the simulated area can be modeled by having mosquitoes migrate to an external node.

Finally, we have largely avoided including fitness costs associated with resistance or mosquitoes carrying drives that could have large effects on the outcomes of gene drive or vector control based intervention strategies in the field. There are likely fitness costs in insecticide-resistant vectors [61] or genetically modified mosquitoes [62]. Additionally, suppression through gene drives can be achieved in a number of ways including fecundity reduction or reduced egg batch size. Sex distortion is another modality of suppression. However, while the framework of the model lends itself to including inheritance logic when a sex distorter drive is used, we currently have not implemented it within the model, and fitness costs will have to be carefully characterized by field or lab data before the model can predict outcomes for different characteristics associated with vector genetics in a particular region.

## Conclusion

An agent-based model of vector genetics that can account for insecticide resistance and gene drives is presented here. When embedded into an agent-based model of malaria immunity and transmission dynamics, it can be used to simulate the evolution of insecticide resistance in a range of transmission settings with ongoing vector control interventions. The impact of insecticide resistance in a high transmission setting with repeated deployments of insecticide-treated nets was simulated as an example. The results suggest periodic exposure over a number of years to the same insecticides can lead to selection of resistant vectors despite low prevalence of resistance in a highly seasonal setting. While the effects of resistance are minor after the first round of exposure to insecticides, subsequent rounds can accelerate resistance, which could lead to rapid resurgences in malaria prevalence and burden.

The model also provides a flexible framework to evaluate expected impact of new tools in programmatic settings. For example, gene drives could be a powerful tool in the fight against malaria. However, gene drives alone may not be able to eliminate malaria. As an example, the vector genetics model presented here was leveraged to simulate a scenario when gene drives are combined with traditional vector control tools such as ITNs in a highly seasonal and high transmission setting. Preliminary results from these simulations suggest a combination of gene drives and traditional vector control methods offer a more robust strategy to achieving malaria elimination than deploying each of these interventions independently.

## Supporting information

The code, input files, and model executable for all simualations can be found on GitHub (https://github.com/InstituteforDiseaseModeling/selvaraj_vector_genetics_2020). Software dependencies such as dtk-tools, dtk-tools-malaria, and the malaria-toolbox packages are publicly available or available upon request from

**S1.**
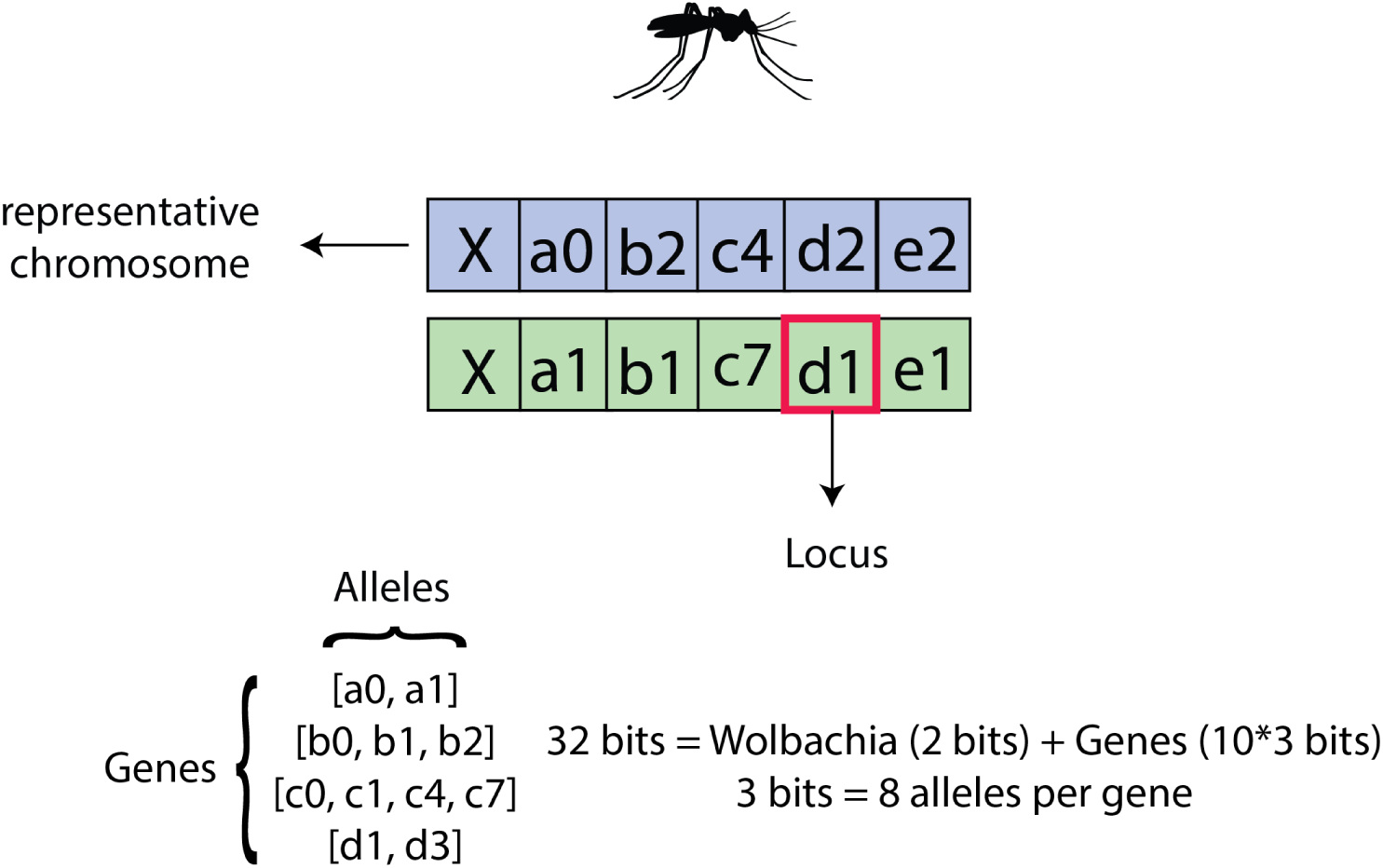
Memory allocation for vector genetics model embedded in EMOD. Each vector or vector cohort carries with it 64 bits of memory dedicated to a diploid carrying 10 different representative genes with up to 8 different alleles per gene. 4 bits are reserved for microbial interventions such as Wolbachia, metrhizium or microsporidia. While a mosquito has 3 chromosome, the representative genome here consists of only one pair of representative chromosomes.

**S2.**
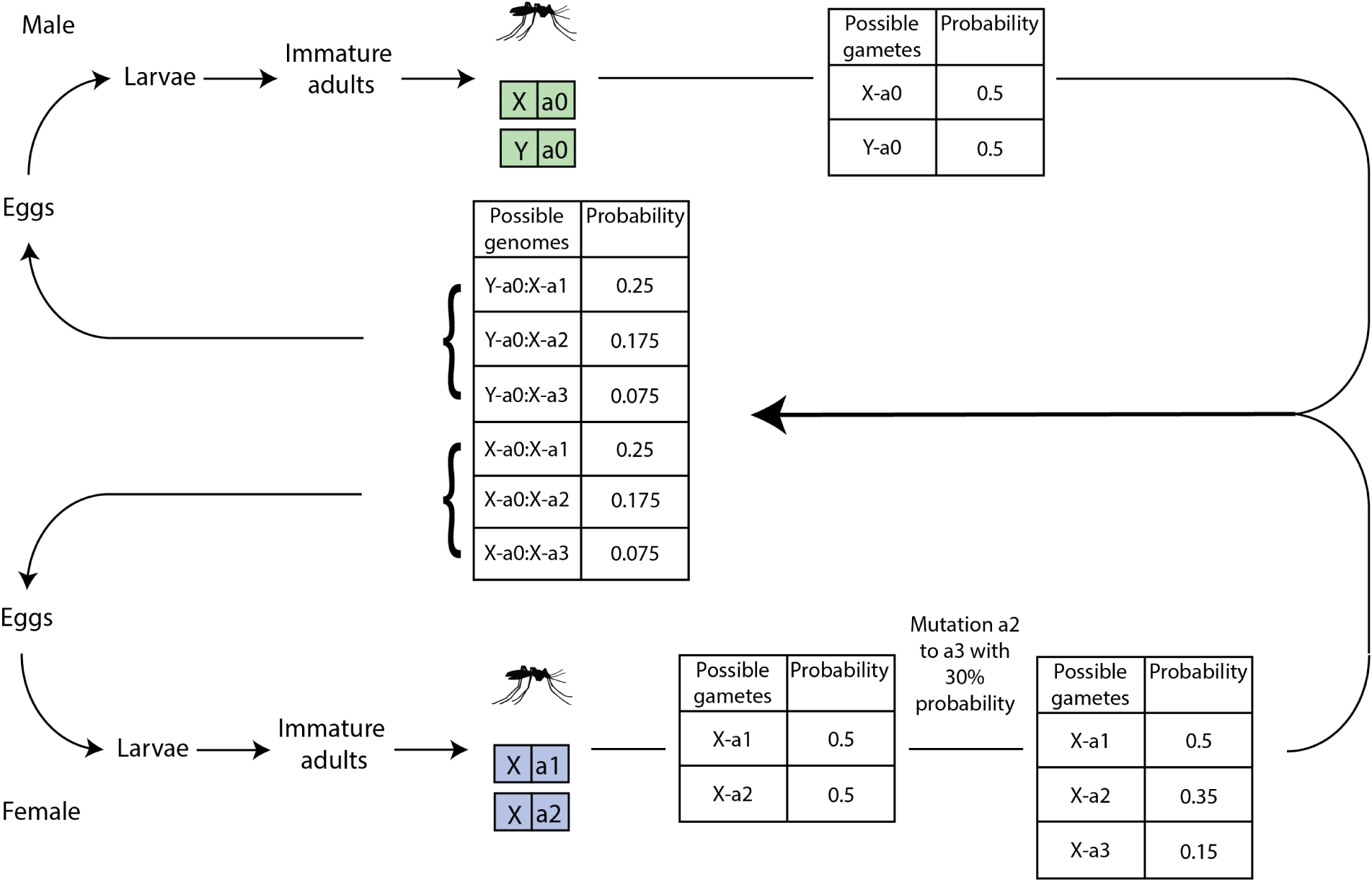
Mendelian inheritance of vector genes in EMOD. EMOD adopts a 4 part lifecycle for the vector starting from eggs that progress to the larval stage before moving onto the immature adult and adult stage. When adults mate, genomes from male and female mosquitoes are used to calculate the likelihood of existence of a gamete carrying a certains set of allele combinations. Random mutation are then applied and possible genome probabilities calculated. Mutation and recombination rates used here are illustrative and do not reflect a specific real world phenomenon. These probabilites are multiplied by the egg batch size, which is modeled as a phenotypic property, for each species to obtain the number of eggs bearing each genome.

**S3.**
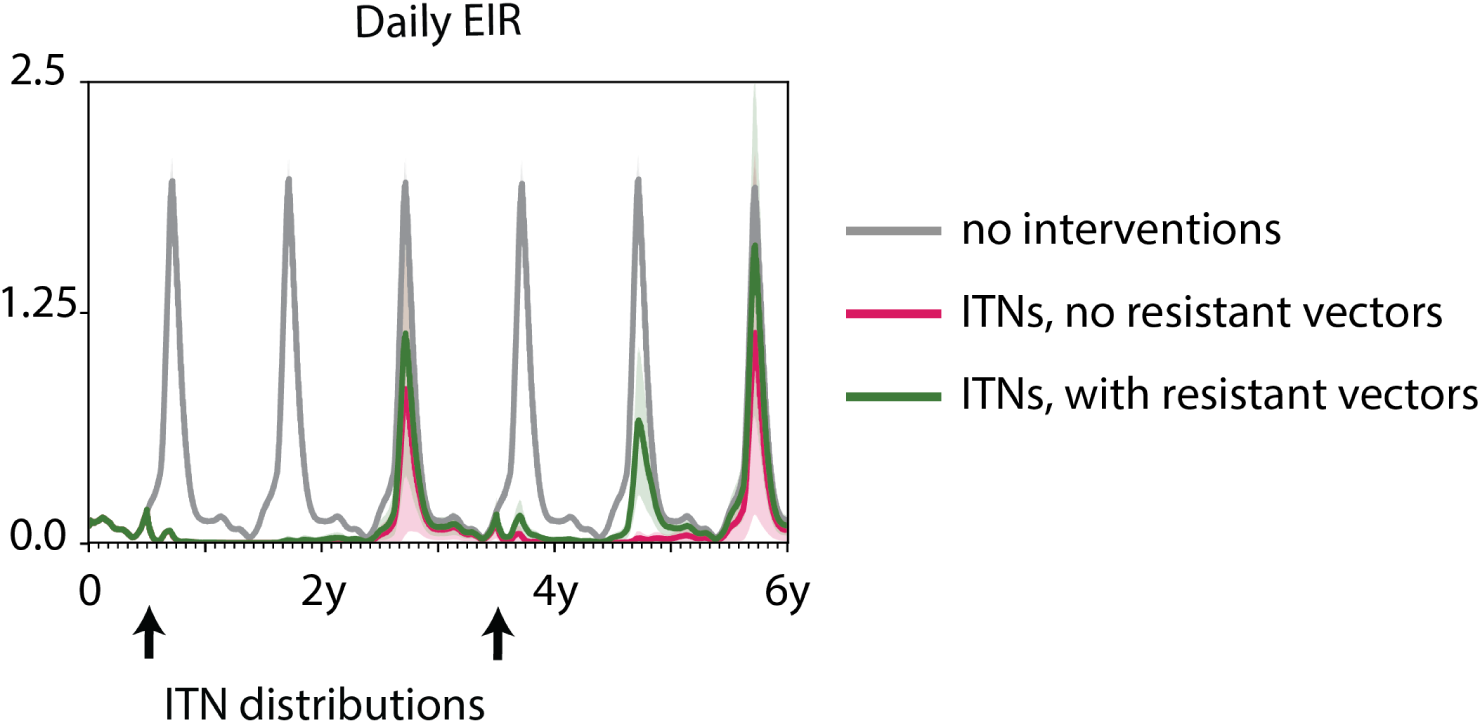
Daily EIR across a period of six years averaged over 50 stochastic realizations for three scenarios – when no interventions are deployed, when ITNs are deployed at 60% coverage absent any resistance, and with the presence of resistance. The shaded area around the mean represents one standard deviation calculated across the 50 stochastic realizations. ITNs are distributed every three years at the beginning of the peak season.

**S4.**
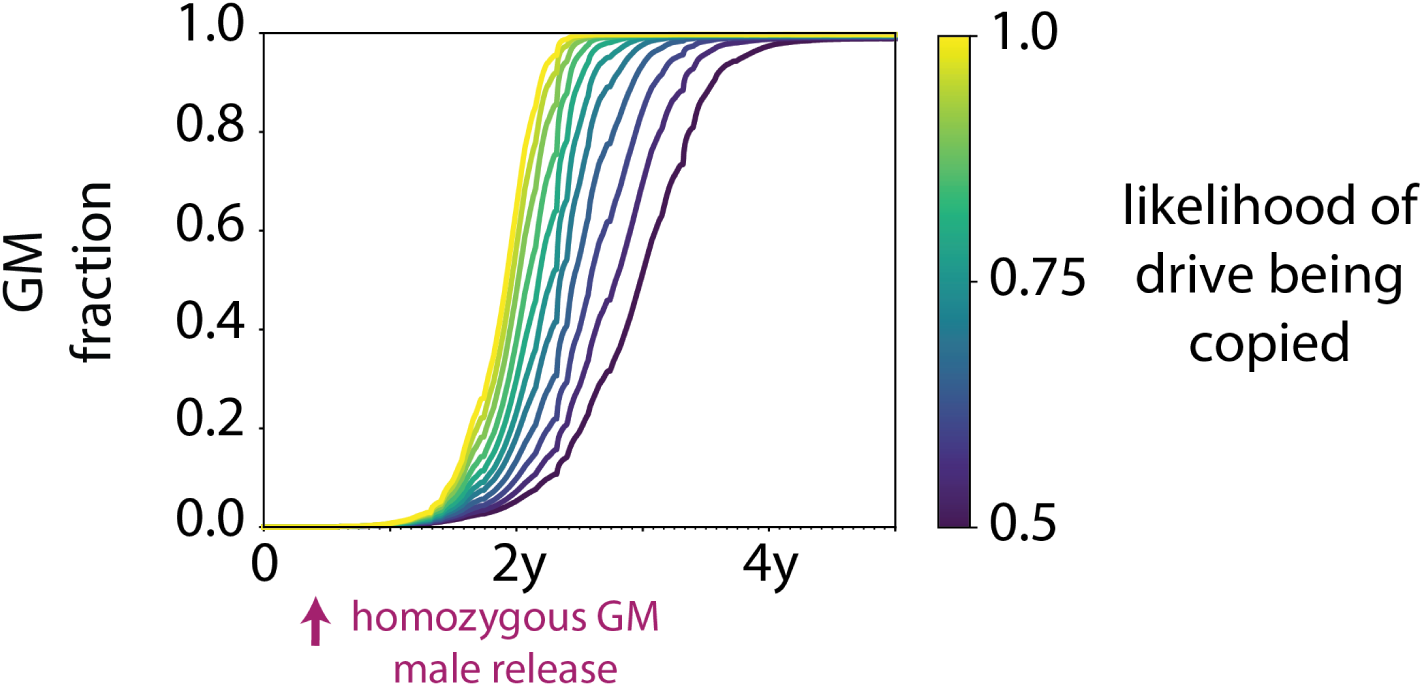
Establishment rates for different probabilities of the drive being copied over to the target location. There are no other vector control interventions. And transmission to human is set to 0.

**S5.**
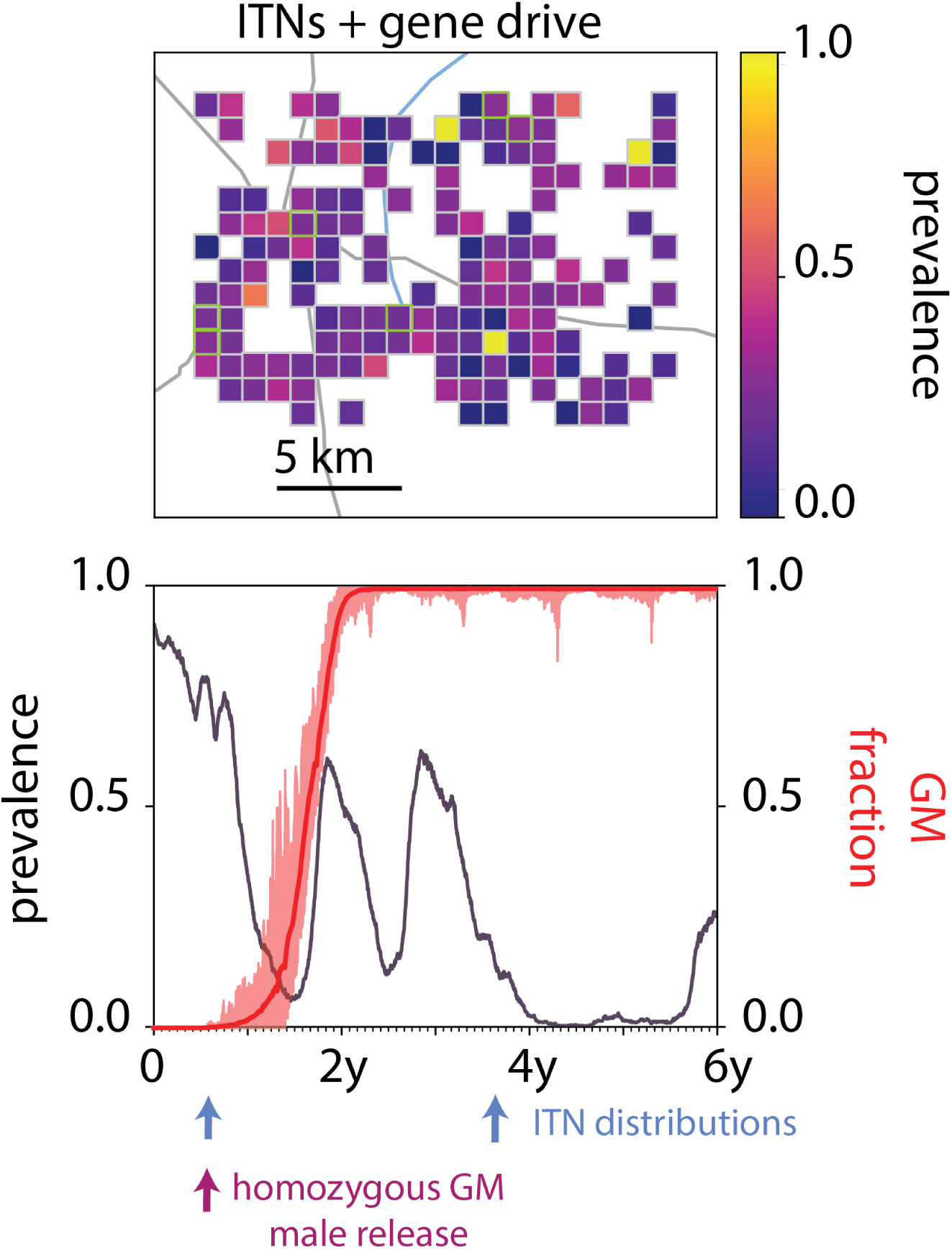
Single realization of the combined ITN and gene drive scenario where elimination was not attained at end of a six year simulation. Upper panel describes spatial distribution of prevalence at the end of six years. Blue line in lower panel describes total prevalence in simulated area. Black line describes overall establishment of GM mosquitoes over time while the red envelope describes the range of establishment rates across all spatial nodes.

## Acknowledgments

The authors would like to thank Austin Burt and Charles Wondji for helpful discussions on gene drives and insecticides, respectively. We would also like to thank Svetlana Titova, Clinton Collins and Jeff Steinkraus for software support.

